# Investigating and Modeling the Factors that Affect Genetic Circuit Performance

**DOI:** 10.1101/2022.05.16.492150

**Authors:** Shai Zilberzwige-Tal, Pedro Fontanarrosa, Darya Bychenko, Yuval Dorfan, Ehud Gazit, Chris J. Myers

## Abstract

Over the past two decades, synthetic biology has yielded ever more complex genetic circuits able to perform sophisticated functions in response to specific signals. Yet, genetic circuits are not immediately transferable to an outside-the-lab setting where their performance is highly compromised. We propose introducing a scale step to the design-build-test workflow to include factors that might contribute to unexpected genetic circuit performance. As a proof-of-concept, we designed and tested a genetic circuit under different temperatures, mediums, inducer concentrations, and bacterial growth phases. We determined that the circuit’s performance is dramatically altered when these factors differ from the optimal lab conditions. Based on these results, a scaling effort, coupled with a learning process, proceeded to generate model predictions for the genetic circuit’s performance under untested conditions, which is currently lacking in synthetic biology application design. As the synthetic biology discipline transitions from proof-of-concept genetic programs to appropriate and safe application implementations, more emphasis on a scale step is needed to ensure correct and robust performances.

## 1 Introduction

Synthetic biology aims to address pressing global challenges including disease diagnosis and treatment [1, 2], bio-fuels production [3, 4, 5], contamination detection [6], bio-manufacturing [7, 8, 9, 4, 10, 11], etc. These are achieved by engineering biological systems with new capabilities, granting cellular control and user-defined performance [12, 4]. The variety of genetic circuit functions include a genetic toggle switch [13], genetic counters [14], low- or high-frequency filters [15, 16], adders [17], sequential asynchronous logic circuits [18], and more.

An implicit, iterative *Design-Build-Test* (DBT) process is often used to develop these ingenious genetic circuits. However, bias is introduced into the DBT process in almost all of its steps and the variability of environmental factors that affect a circuits’ behavior is often not taken into account. This might hinder a circuit’s expected performance when applied outside-the-lab (OOT). Models used by *genetic design automation* (GDA) tools are mostly based on experiments carried out under optimal lab conditions (OLC) [19, 20, 21, 22]. Furthermore, most rely only on the expression of a fluorescent protein as an output reporter under OLC. This setup leads to an inaccurate *Scale* step with regard to the actual circuits’ performance when applied in non-OLC that can produce erroneous or faulty behavior with unpredictable outcomes. Furthermore, with a narrow *Test* step, the *learning* usually is limited to a post-hoc description of circuit dynamics. This would be especially perilous for engineered systems that are aimed to operate in dynamic environments, such as living therapeutics and whole cell biosensors.

This study applies a broader *Test* step to a designed delay-signal circuit to include more environmental dynamic factors and reporters (as shown in Fig. 1). The circuit’s output, as well as the time for output detection, were observed to be highly variable for different temperatures, mediums, inducer concentration, bacterial growth-phases, and output reporters. If the performance of the delay circuit is compromised by the tested experimental factors presented here, it will inevitably alter its behavior in other contexts, which would not have been predicted by GDA tools.

**Figure 1:**
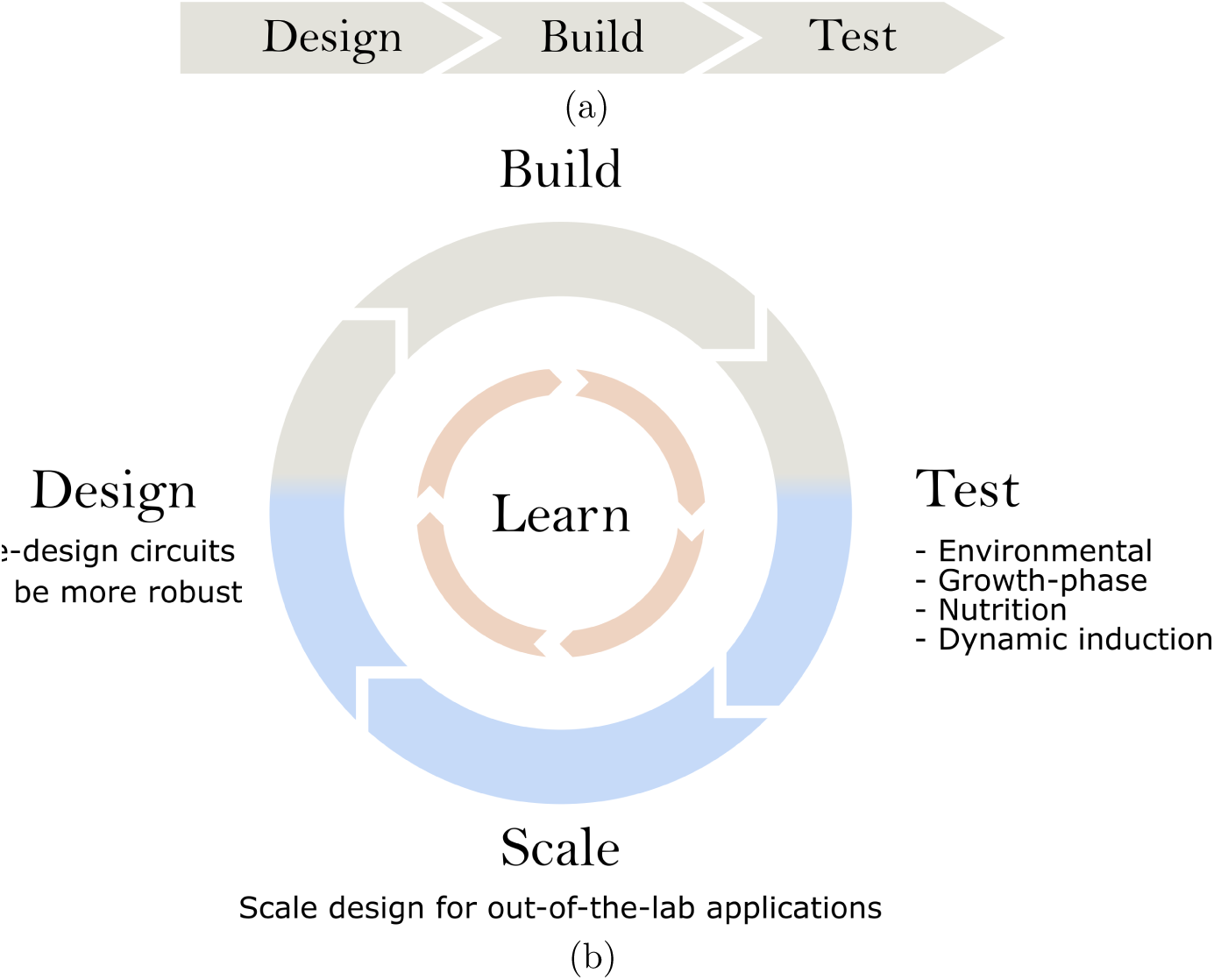
DBTS workflows in synthetic biology. **a** Most common workflow, where testing is done only in OLC and there is no feedback to alter design. **b** Proposed workflow in which the *Test* step includes different conditions that may affect the outcome of a circuit. This can be utilized in the *Scale* step to obtain new model predictions, which will allow one to make better design choices. Ultimately, this will produce robust designs.

We propose to introduce a *Scale* step as part of a new and improved Design-Build-Test-Scale (DBTS) process. Scaling refers to the process of considering the variability of factors that can affect genetic circuit performance in real-life applications. Most studies either have a non-existent *Scale* step, or it consists only of a post-hoc description of the designed circuit’s performance at OLC. This work not only provides a re-parametrization effort for different experimental conditions, but also produces a new model to determine the necessary predictions for untested conditions. As a case study, we focused on the effect of growth phase on the circuit’s output, in which we observed a trend in delay and total output production. This, in turn, allowed for a deeper *Scale* step which ultimately resulted in a new model that estimates these trends, thus enabling the capacity to predict untested delays and output production of the circuit which can be further applied for scaling.

Thus, we propose that a greater emphasis in the *Test* and *Scale* steps of a DBTS cycle are needed to build more predictive models and to reduce bias across the entire DBTS cycle. This, in turn, will enable the possibility of finding design alternatives to any unexpected behavior and performance when the circuits are used in applications, improving a genetic circuit’s robustness [23]. As we move from proof-of-concept designs to more real-life applications, a thorough *Test* step provides the necessary data that allows for a significant *Learn* step and, therefore, an appropriate *Scale* step.

## 2 Results

### 2.1 Designed Circuit and Predicted Behavior

Fig. 2a shows a schematic of the actual circuit designed, built, and tested in this work. For more information on the layout design, please refer to the Methods.

**Figure 2:**
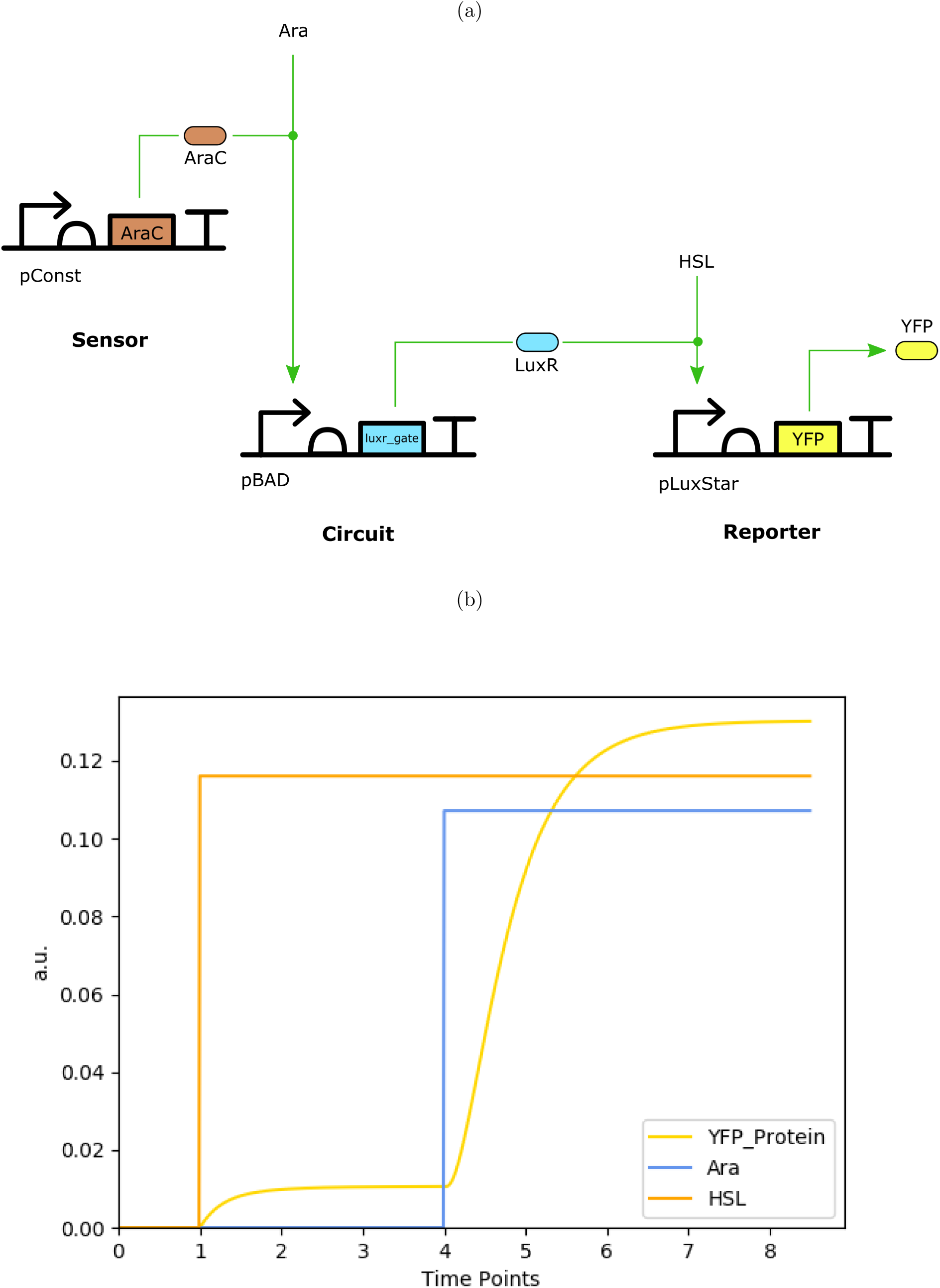
(a) Designed delay circuit using Cello gates [20]. Sequences obtained from SynBioHub [24] ^1^. This circuit produces *yellow fluorescent protein* (YFP) after a delay when both *arabinose* and *oxohexanoyl-homoserine lactone* (HSL) are present. (b) Delay circuit simulation results using default parameters. YFP production (in a.u.) over increasing simulated time-points.

The intended purpose of this design is to provide some delay between an input concentration change and output production, whilst avoiding unwanted switching behavior due to initial propagation of an erroneous state. The circuit will not produce *yellow fluorescent protein* (YFP), unless both *arabinose* (Ara) and *oxohexanoyl-homoserine lactone* (HSL) are present. This design avoids unwanted production of an output (YFP) when the circuit is initialized in a cell without Ara: even if there is initial production of LuxR. Meaning, if there is no HSL, the circuit will not produce YFP and both Ara and HSL are needed to produce the circuit’s output.

The initial model predictions of the circuit, shown in Fig. 2b, were done in iBioSim [25] using an automatic model generator to produce an *ordinary differential equation* (ODE) model of the circuit. The resulting complete model was then analyzed using the Runge-Kutta-Fehlberg method [26], also implemented in iBioSim [25].

The simulation results show that there is no YFP production when only HSL is present and, furthermore, there is a delay in the YFP production when Ara is added as expected. However, given that these simulations are using default parameters that were characterized under OLC (obtained from [20]), these only provide qualitative information on how the actual circuit is going to behave only when tested in OLC.

### 2.2 Control Experiment

Using iBioSim [25] and standard genetic parts [20], the delay circuit was designed aiming for a relative output time delay post-induction. To test the actual delay, a simple control experiment was set using OLC. Briefly, bacteria were cultivated in M9 glucose media at 37°C in the presence of both inducers from T=0 (Ara and HSL), which simulate the characterization assays that were done in Cello [20] (Fig. 3 a). As negative controls, the bacteria were also cultivated without any inducer or with the presence of only one of the inducers. Under these conditions, it took an average of 180 minutes to detect a fluorescence signal (Fig. 3 b(i) and b(ii)). The 180 minutes was set as the *optimal detection time* (ODT) and the signal intensity from this assay was also set as the *optimal intensity* (OI) since both were measured under optimal growth conditions.

**Figure 3:**
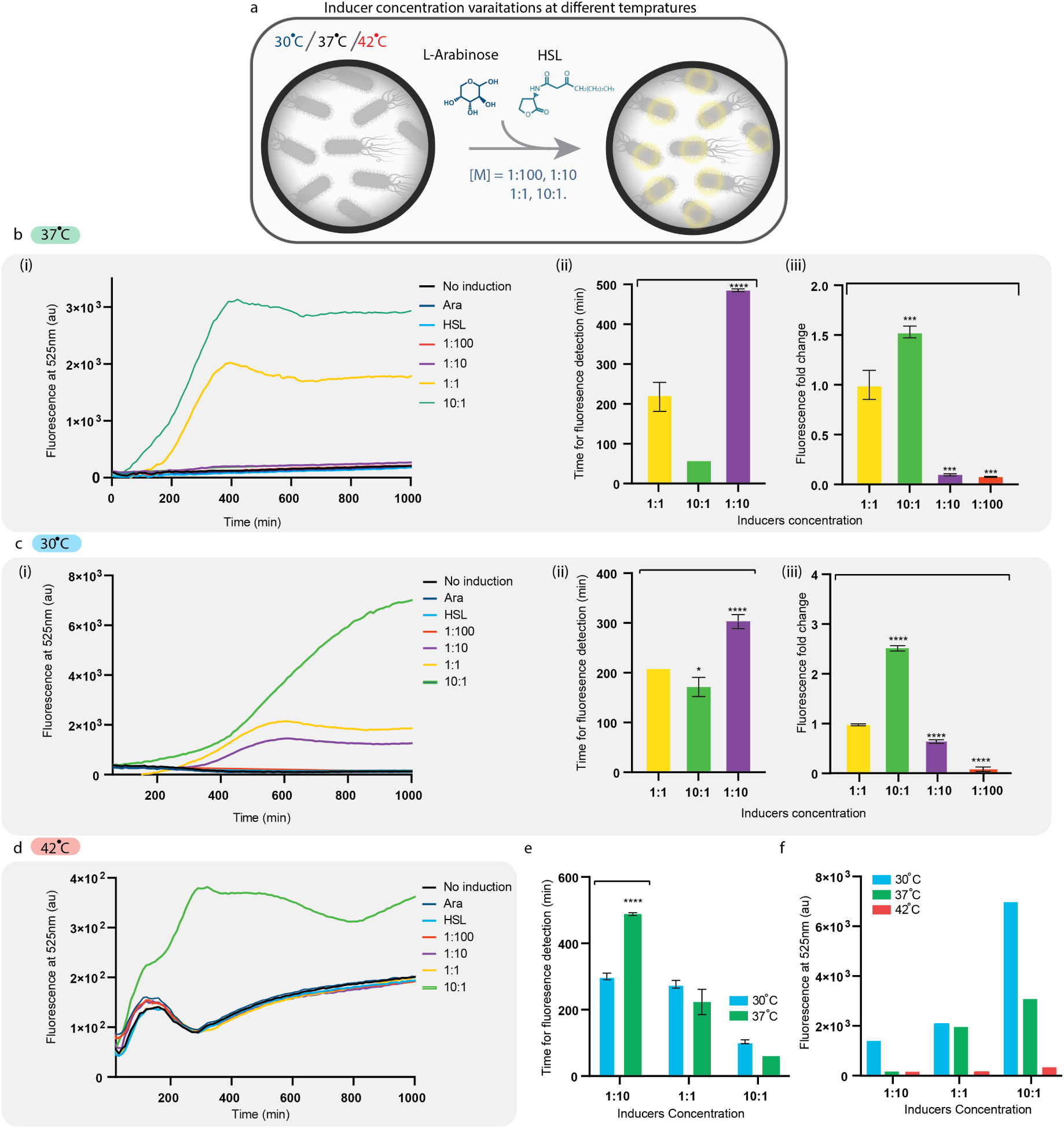
Control assay and inducer concentration variations at different temperatures. **a** Scheme of the control and inducers’ concentration variations assays. Ranging inducer concentrations were added at T=0, 37°C, 30°C and 42°C. **b** Measured fluorescence signal (in a.u.) over time **(i)** of the control assay and of bacteria that were cultivated with a range of inducers concentrations. **b(ii)** Comparison of the time to detect a fluorescence signal from bacteria cultivated with different inducers concentrations. \**P* < 0.01, \*\*\**P* < 0.005 (Student’s *t*-test). **b(iii)** Comparison of the maximum fluorescence signal intensity fold change detected from bacteria that were cultivated with different inducers concentrations. \*\*\**P* < 0.005 (Student’s *t*-test). **c(i)** Measured fluorescence signal (in a.u.) over time of bacteria cultivated with different inducers concentrations at 30°C. **c(ii)** Comparison of the time to detect a fluorescence signal. \**P* < 0.01. \*\*\**P* < 0.005 (Student’s *t*-test). **c(iii)** Comparison of the maximum fluorescence signal intensity. \*\*\**P* < 0.005, \*\*\*\**P* < 0.001 (Student’s *t*-test). **d** Measured fluorescence signal (in a.u.) over time of bacteria cultivated with different inducers concentrations at 42°C. The differences between the negative controls and the induced samples were not significant and therefore cannot be plotted in fluorescence detection and fold change graphs. **e** Comparison of the time to detect a fluorescence signal at 30°C and 37°C from bacteria cultivated with different inducer’s concentrations. **f** Comparison of maximum fluorescence signal at 30°C, 37°C and 42°C from bacteria cultivated with different inducer’s concentrations.

Next, the circuit’s robustness was tested at different conditions that mimic the dynamic environments that bacteria may encounter OOT. The following sections will describe the genetic circuit’s dynamic performance under different testing conditions. The purpose of these testing conditions is to observe how it will affect the time for signal detection and signal intensity.

### 2.3 Inducer Concentrations

An important dynamic variation bacteria could encounter when applied OOT is inducer concentrations. The inducers concentrations that were set in our control experiment as 1:1 are 2mM and 2*μ*M for Ara and HSL, respectively, since these are the concentrations that were used in Cello [20]. The bacteria were cultivated in the presence of serially diluted concentrations (1:100 and 1:100) and in concentrated concentration (10:1) (Fig. 3a). Bacteria that were cultivated in the presence of the 10:1 inducers, were able to produce a fluorescence signal much faster than the ODT (Fig. 3b(i) and b(ii)). In addition, the signal intensity of the 10:1 sample was significantly higher than the signal from OI (Fig. 3b(iii)). However, in the lower concentrations of 1:100 and 1:10, the fluorescence signal was weak and barely detected using our methods (Fig. 3b(i) and b(iii)). Thus, the genetic circuit behavior in terms of time for signal detection and its intensity is highly compromised and dependent on the inducer concentrations. Thus, the genetic circuit behavior is not robust in terms of time for signal detection and signal intensity.

### 2.4 Temperature

Among environmental conditions, temperature is an important, if not the most important, variable affecting microbial growth and as a direct consequence we hypothesize that it affects the genetic circuit’s behavior [27]. *E. coli* can survive in a range of temperatures starting from 4°C up to 45°C, with an optimal growth at 37°C, and it is usually the temperature used for cultivating *E. coli* in the lab [28]. *E. coli* was cultivated at 30°C and 42°C in the presence of a range of inducer’s concentrations as described above (Fig. 3a). Lowering the growth temperature from 37°C to 30°C resulted in a higher fluorescence signal of both 10:1 and 1:10 inducers’ concentrations (Fig. 3 f). It should be noted that in addition to a lower fluorescence signal of the 1:10 dilution at 37°C, the time to detect the signal was significantly higher (Fig. 3e). The circuit’s behavior was tested at 42°C in the presence of different inducers’ concentration as mentioned above. At this temperature, a fluorescence signal was observed only in the high inducers’ concentration (10:1) (Fig. 3d) and the intensity of the detected signal was an order of magnitude lower than the OI (Fig. 3f). We assume that the burden of the genetic circuit on the bacteria is higher at 42°C than 37°C, since in the higher temperature the bacteria is in greater stress due to increased expression of heat shock proteins, which add to the overall burden on the cell [29]. Thus, the behavior of the circuit at the higher temperature differ significantly from its behavior during the control assay at 37°C.

### 2.5 Soil

Medium composition can be highly diversified, which may lead to altered genetic circuit’s behavior. We decided to test our genetic circuit in a medium that will mimic applications OOT. Therefore, bacteria were cultivated in sterile and non-sterile soil (Fig. 4a). A fluorescence signal was detected in both samples, however the signal intensity was higher in the sterile soil sample (Fig. 4b and c). This can be attributed to the fact that in the sterile sample there is less competition for nutrients, since the sample was autoclaved. While in the non-sterile sample, there are many other bacteria competing for nutrients, which leads to a lower signal. It should be noted that the production of the fluorescence signal in the soil samples was maintained for a longer period of time than in all other conditions (Fig. 4). This, again, can be attributed to the fact that the soil contains nutrients that are lacking in our control assay medium (see Methods).

**Figure 4:**
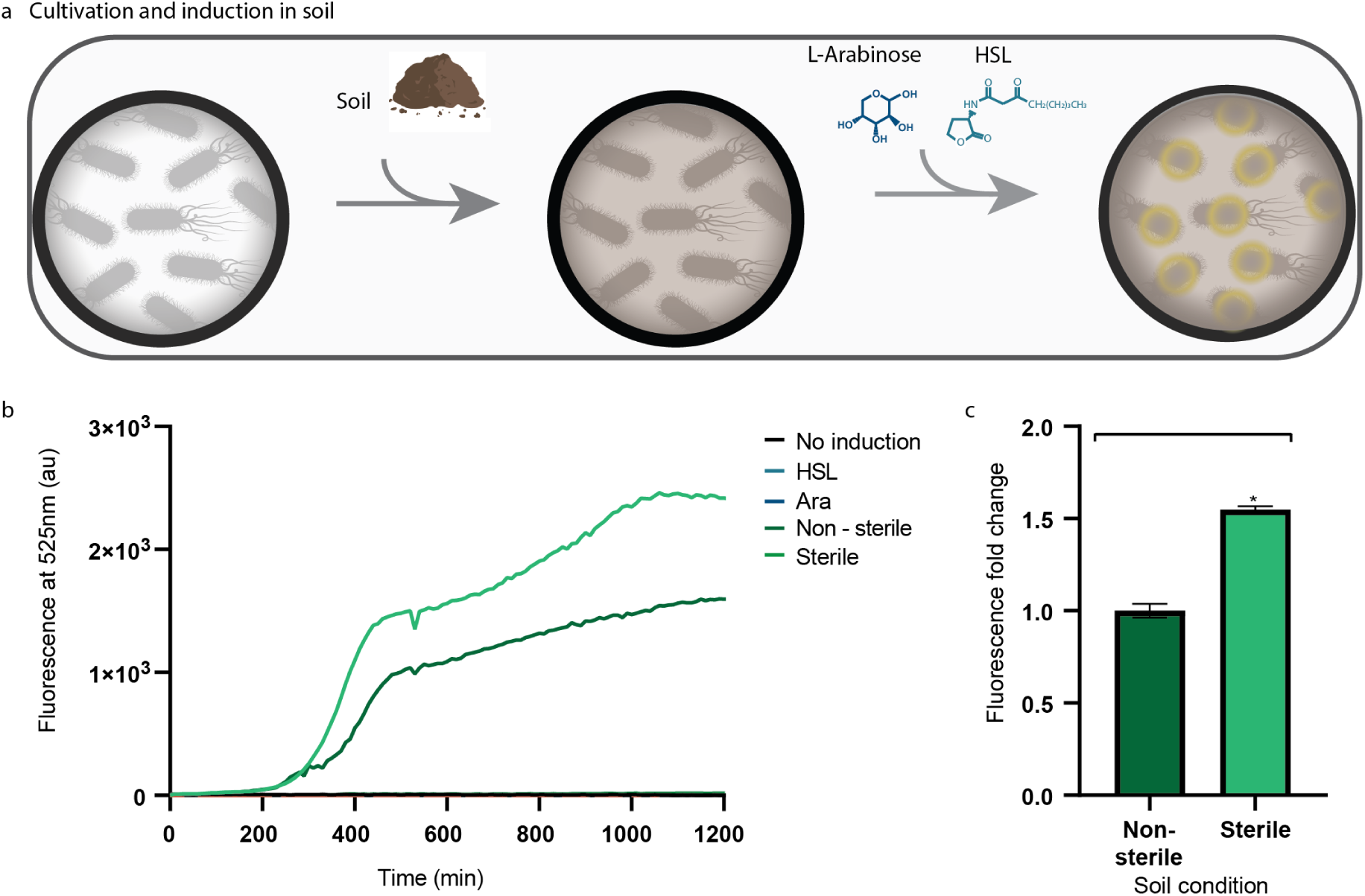
Cultivation in media containing 2% sterile and non-sterile soil. **a** Scheme of the cultivation with soil assay, 2% (W/V) soil was added to the media and then both HSL and Ara were added at T=0, which induced the production of YFP. **b** Measured fluorescence signal (in a.u.) over time of the bacteria cultivated in 2% (W/V) sterile and non-sterile soil. **c** Comparison of the maximum fluorescence signal intensity detected from bacteria that were cultivated in 2% sterile and non-sterile soil. \**P* < 0.01. (Student’s *t*-test)

### 2.6 Induction at Different Bacterial Growth Phases

Optical density (OD) measurements are usually used for quantifying the growth of a bacterial culture [30]. When the data is plotted semi-logarithmically, four growth phases are distinguishable: (i) lag phase which is non-replicative; (ii) exponential phase which is replicative; (iii) stationary phase where growth ceases, but cells remain metabolically active and (iv) death phase where there is a gradual decline in viable cells. During these phases, the transcriptome and proteome of the bacteria dramatically change, as well as the transcription and translation rates. To test the robustness of the time delay circuit across the different growth phases, we grow the bacteria in the presence of HSL from T=0, and we added the second inducer, Ara, at different growth phases: early-lag (T=0, such as the control experiment), late-lag, early-exponential, mid-exponential, late-exponential and stationary, and measured the fluorescence signal over time (Fig. 5a). When the inducers were added at a later phase than the early-lag (*EL*), which is also the ODT, the time for fluorescence detection was significantly decreased (Fig. 5b and c). Interestingly, we observed that the decrease was gradual from the late-lag (*LL*) to early-exponential (*EE)* and middle-exponential (*ME)*. This was followed by a minor increase from late-exponential (*LE)* and stationary (*S)* (Fig. 5c). When comparing the signal intensity of the different induction times, it is clearly shown that the signal decreases when induction starts at a later stage, especially at *ME, LE*, and *S* (Fig. 5b and d). It should be noted that the encounter with the inducer at different growth phases did not influence the doubling time of the bacteria when compared to the doubling time of the bacteria without induction (Supplementary Fig. 1. The behavior of the time-delay genetic circuit alters when the bacteria are induced at different growth phases, suggesting that the unsteadiness of the bacteria transcription and translation rates influences the behavior of the genetic circuit.

**Figure 5:**
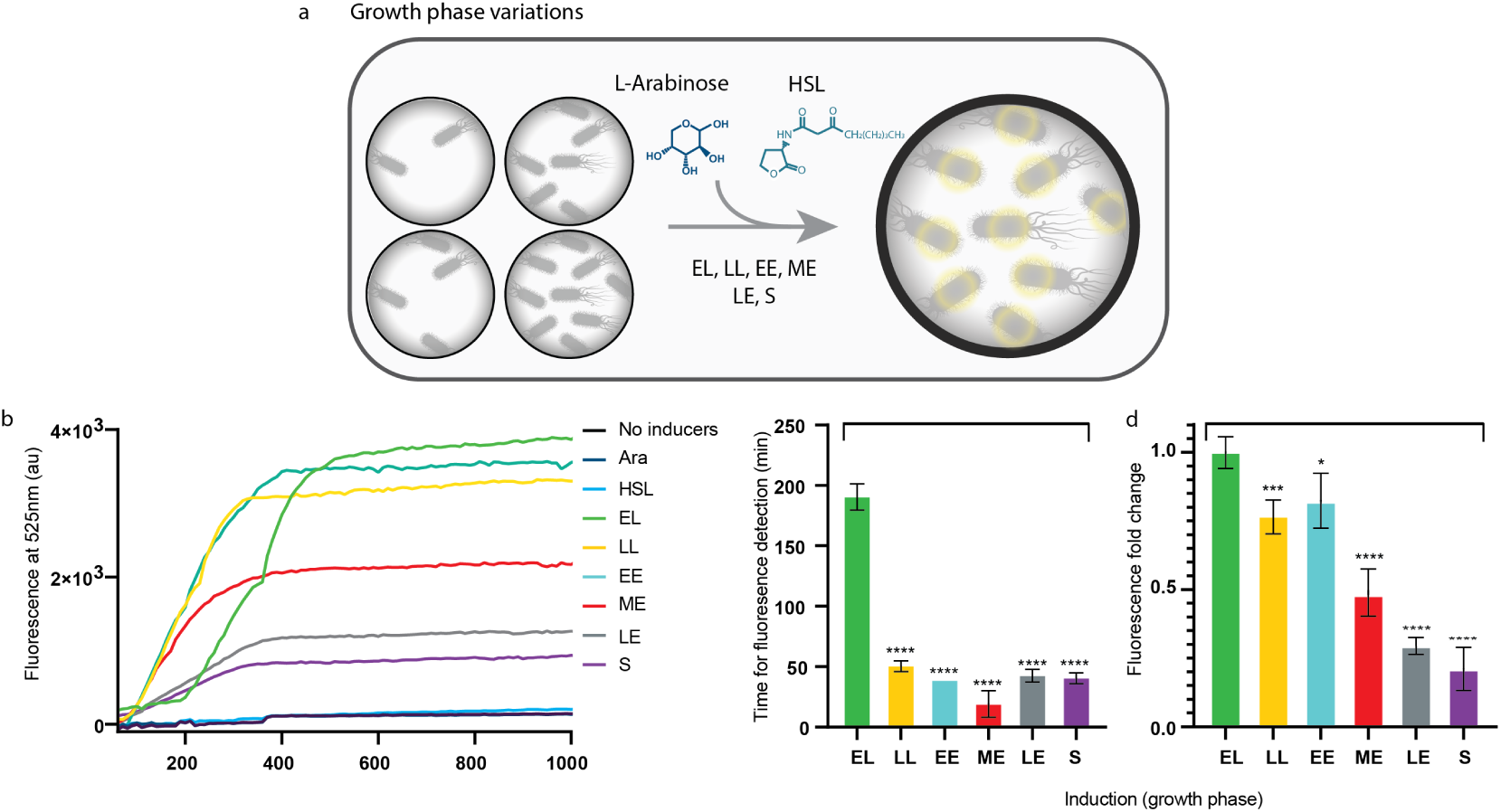
Growth phase variations assay. **a** Scheme of the growth phase variations. HSL was added at T=0 (EL) and Ara was later on added at a different growth phase which induced the production of YFP. **b** Measured fluorescence signal (in a.u.) over time of bacteria induced at different growth phases. **c** Comparison of the time to detect a fluorescence signal from the bacteria. \*\*\*\**P* < 0.001 (Student’s *t*-test). **d** Comparison of the maximum fluorescence signal intensity fold change detected from the bacteria. \**P* < 0.01, \*\*\**P* < 0.005, \*\*\*\**P* < 0.001 (Student’s *t*-test).

### 2.7 Total Fluorescence Production

To address the different genetic circuit behavior when induced at different growth phases, we calculated the total amount of fluorescence in *arbitrary units* (a.u.) that are produced hourly following the detection of the fluorescence signal (see Methods). Our hypothesis is that genetic circuits that are induced during the exponential phase, in which bacteria are dividing rapidly and thus producing more proteins, the hourly a.u. production rate will be higher than genetic circuits that are induced in the lag or stationary phase, where protein synthesis is much lower [30]. Moreover, genetic circuits that are induced at a later growth phase will not produce as much total fluorescence a.u. as genetic circuits that are induced earlier, since the bacteria are closer to the stationary phase and start to die due to the lack of nutrients. Thus, we decided to compare the accumulated fluorescence a.u. production in bacteria that are induced at different growth phases (Fig. 6a). According to the results, in the first few hours, the highest fluorescence a.u. production was of the LL and EE induction (Fig. 6b). However, following four hours, the hourly fluorescence a.u. production of the LL, EE and ME were relatively similar (Fig. 6b). Through all the first five hours, the LE and S hourly fluorescence a.u. production were the lowest (Fig. 6b). These findings are consistent with our hypothesis regarding the susceptibility of the bacteria to efficiently produce a signal if the induction occurs in an early growth phase. Moreover, the sum of the hourly productions further emphasize that bacteria that are induced at a later stage (starting from the ME phase) are more likely to produce less signal overall (Fig. 6c). Thus, the range of the circuit behavior is heavily influenced by the specific growth phase in which the bacteria are at when they encounter the signal molecules. We hypothesize that the growth phase status is an immensely important factor with implications on protein production and degradation rates, which directly affect gate production rates. This means that growth phase status will affect the parameter values used in the model and, therefore, the predicted behavior. We decided to characterize the gate production rates at different growth phases (see below).

**Figure 6:**
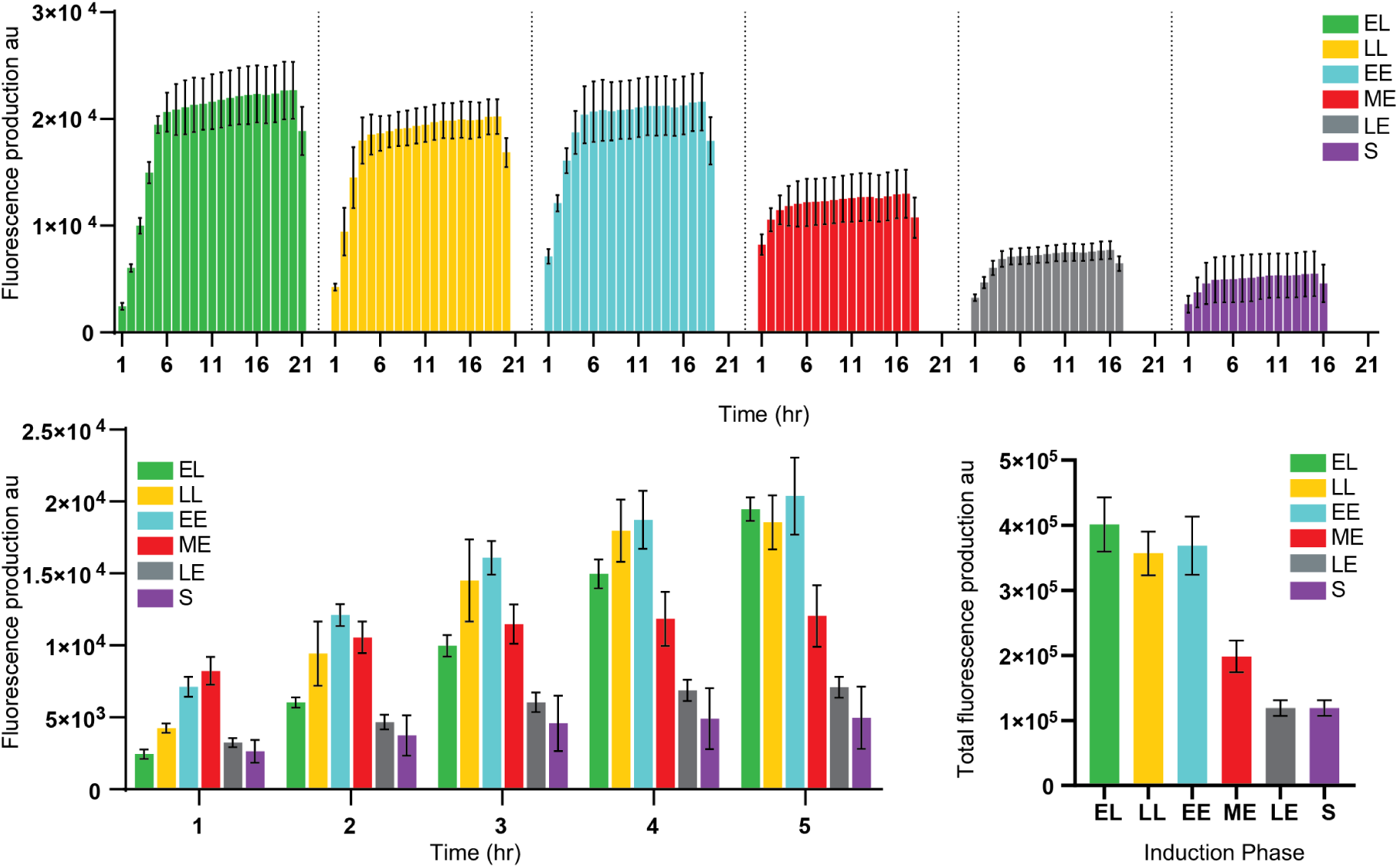
Accumulating fluorescence a.u. production of bacteria induced at different growth phases. **a** hourly accumulated fluorescence a.u. of bacteria induced at different growth phases. **b** Comparison of the hourly accumulated fluorescence a.u. of the first five hours. **c** Comparison of the total accumulated fluorescence a.u. \*\*\*\**P* < 0.001 (Student’s *t*-test).

### 2.8 Gate Re-parametrization

To capture the effect of different growth phases on the production delays, as well as signal intensities of the designed circuit in this work, characterization experiments were done for the different growth phases. These experiments were performed following the methods described in Shin et al. [31] (see Methods). From these experiments, new parameter values for each growth phase were achieved as described below.

#### 2.8.1 Hill Function Parameters

Equation 7 is an equation derived from the Hill function, which is used to calculate the steady-state output of a gate [20, 31]. To acquire the Hill function parameter values used in this equation, gate induction experiments were performed and then fitted as shown in the *“Parametrization Methods”* section. This fitting can be done by minimizing the error of Equation 1 (for *activation*) and 2 (for *repression*) predictions by using parameter fit values and the experimental measurements for the different growth phases as shown in the *“Circuit Induction and Measurements”* section.

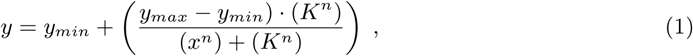

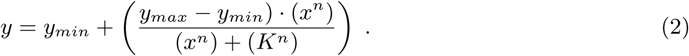

The Hill-function fitted parameter values obtained are shown in Table 3. These parameters can be used to calculate the *steady-state* output of the circuit using Equations 3 and 4. However, to be able to predict the *dynamical* behavior of these circuits, the *τ*_*ON*_ and *τ*_*OFF*_ parameters also have need to be calculated.

#### 2.8.2 Tau (*τ*) Parameters

Equation 8 describes the dynamical response of each gate, using the *τ*_*ON*_ and *τ*_*OFF*_ parameters. To obtain these dynamical parameter values, different *ON-to-OFF* and *OFF-to-ON* characterization experiments were performed using the same gate plasmids and methods as shown in Shin et al. [31] and this work (see Methods). These experiments can be used to fit the data using Equations 3 and 4, obtained from the supplementary material in Shin et al. [31], to achieve parameter values for these conditions. The characterization method used to re-parameterize the dynamical *τ*^*ON*^ and *τ*^*OFF*^ is explained with detail in the Methods section. The results obtained are shown in Table 4.

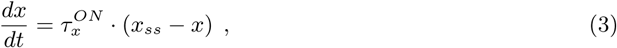

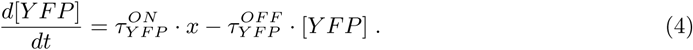

Using the parameters shown in Table 4, derived from the fitting algorithm, new simulations were produced for each growth phase (Fig. 7). The new model simulations predict both lower production of YFP protein (signal intensity), as well as the decrease in time for reaching steady-state, for each successive growth phase, as observed experimentally.

**Figure 7:**
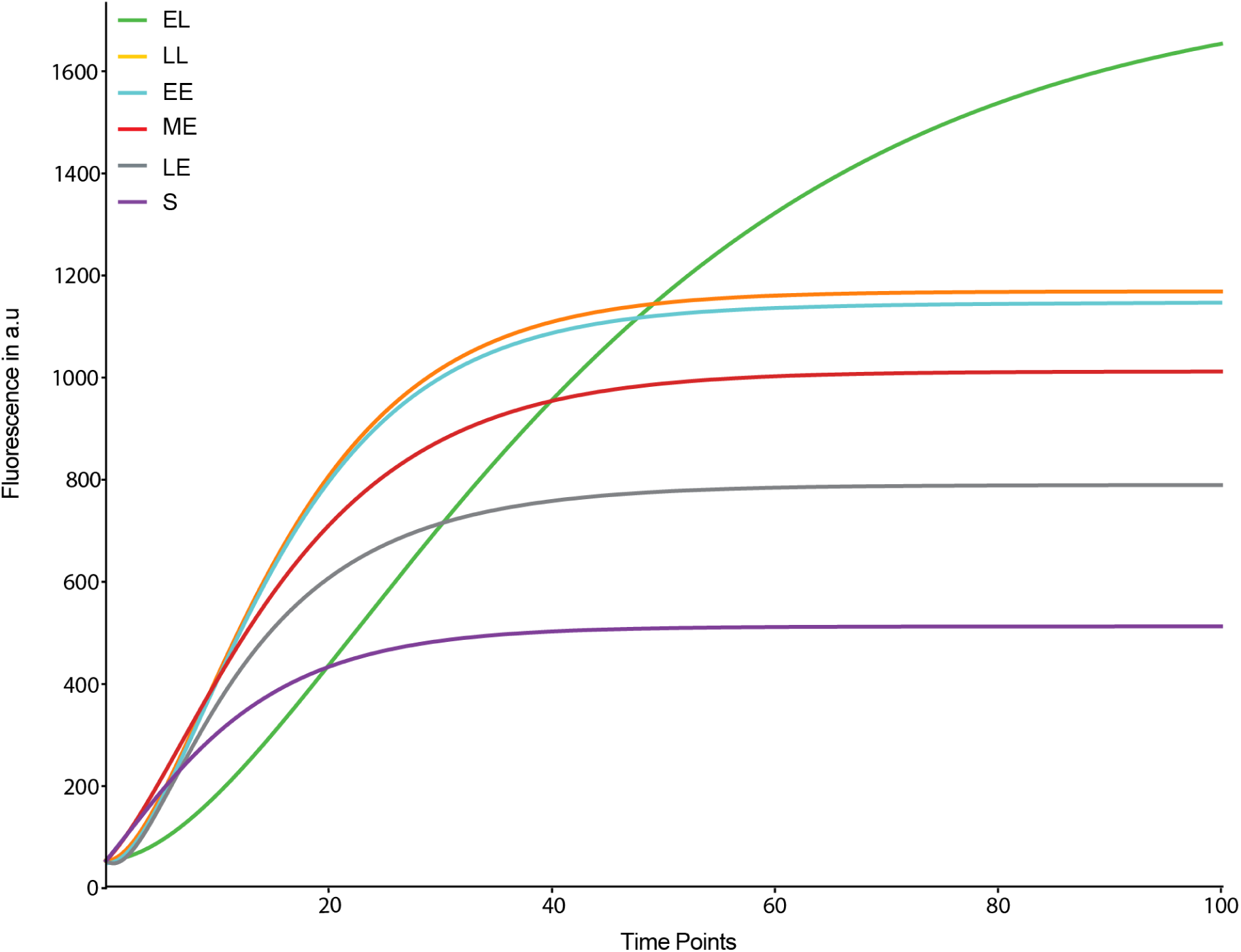
Simulation results for the different growth phases of the AraC gate, using fitted parameters shown in Table 4, obtained using the lmfit Python package [32].

However, the parametrization results (shown in Table 4) were produced without fixing any of the parameter values when using the fitting algorithm, meaning they were all treated as free variables. When there are no fixed parameters, then the fitting algorithm will find the best fit by manipulating the parameter values until the expected outcome best matches the experimental results. This will result in widely different parameter values to compensate for other parameters’ minimization of error while fitting (see for example, *x*_*SS*_ values for different growth phases in Table 4). This means it will be hard to derive any parameter value trends indicative of what might be happening to their magnitude as the circuit is induced at different growth phases.

To discern if there are any parameter value trends that can be attained from fitting the experimental results, we proceeded to fix parameter values to reduce the number of free variables in the fitting algorithm. The first parameter value fixed was 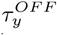. This was done using the fitting algorithm with the ON-to-OFF gate characterization experiments (see *methods* section). Re-fitting the model to the experimental results, with 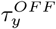 as a fixed parameter value and the rest as free-variables, new parameter values were acquired. These re-fitting iterations were done then by fixing subsequent parameters values by calculating averages from previous iterations. First, *x*_*SS*_ average values were calculated, then fixed for the next iteration, then followed by 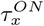. With the last iteration of this re-fitting process, 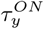 was left as a free-variable, while the rest of the parameters were fixed. This process was carried out to understand the effect of the different experimental conditions on the value of 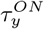. First, since 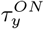 is the parameter closest to the measured parameters in the experiments (YFP fluorescence); and secondly, because if the other parameter values are not fixed, then if there is any parameter value trends, it is lost in the minimization process, when the fitting algorithm tries to find the solutions by increasing a parameter value and decreasing another one. Fig. 8a shows 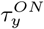 parameter values obtained following this procedure.

**Figure 8:**
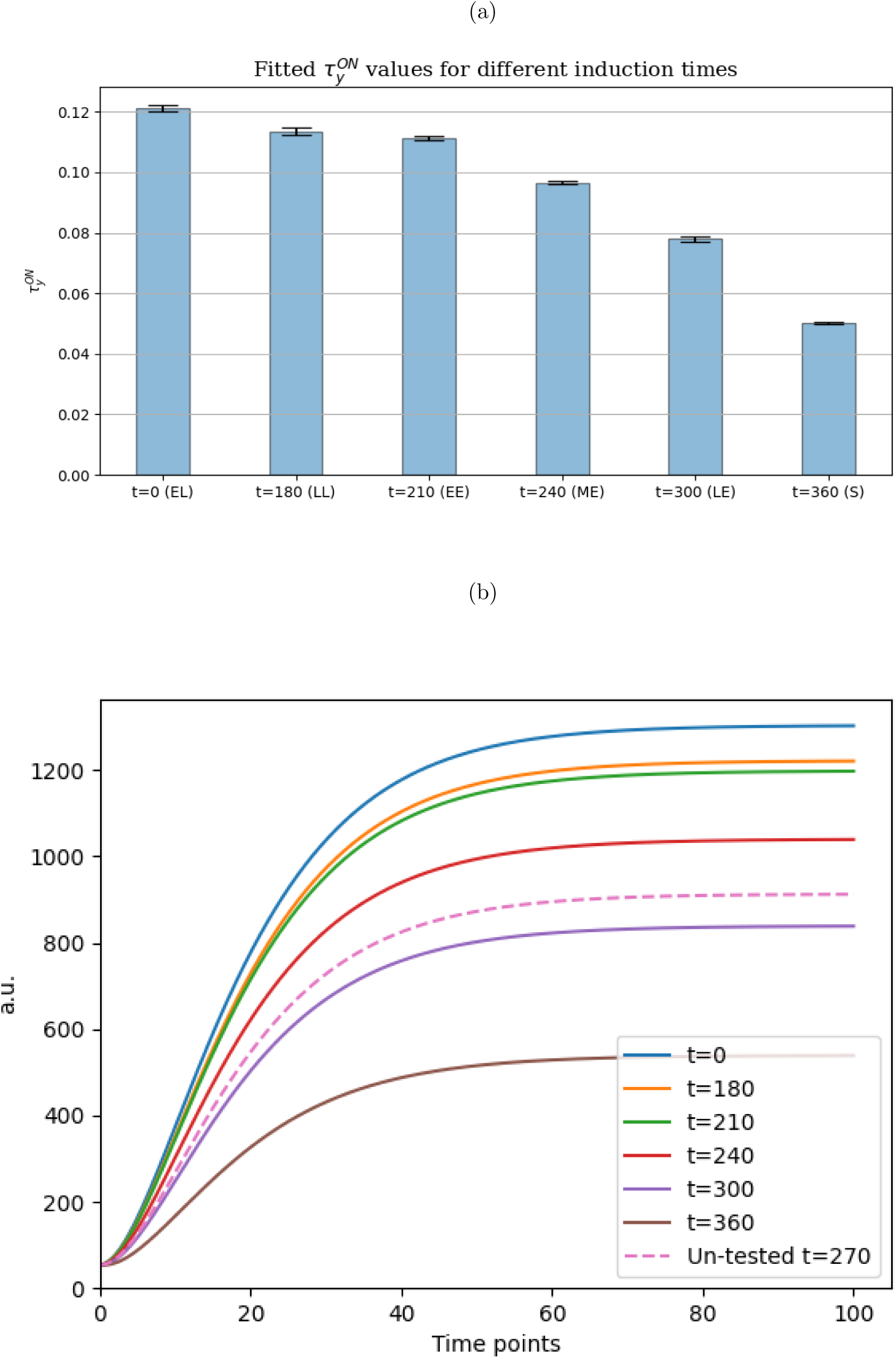
(a) Fitted 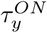 values obtained when fitting fluorescence values for different induction times (t=0, t=180, t=210, etc.) using fitted parameters obtained using the lmfit Python package [32]. (b) Un-tested YFP output prediction for induction time of 270 minutes obtained using the linear regression model to estimate 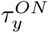’s value for the untested condition.

As hypothesized, the values of 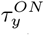 shown in Fig. 8a vary for the different growth phases of the clonal bacteria. When bacteria are induced at a later growth-phase, the values of 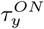 decrease. This coincides with previously observed experimental results where induction at later growth phases decreases the time it took to reach steady-state, as well as the maximum signal intensity at the steady-state. A predictive linear model of signal intensity over time can be created as shown in Equation 5:

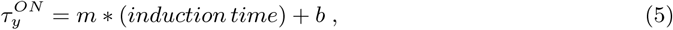

where 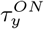 is the predicted value of gate dynamics when the circuit evolves to an *ON* state [31]. Using linear regression, *m* was estimated to be −1.869*e*^*−*04^ and *b* to be 0.13527606. With this equation, a researcher could estimate the value of 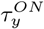 for different times of induction and, therefore, estimate the decrease in output production and delay of a delay circuit for untested conditions as shown in Fig. 8b.

### 2.9 Lysis

The ability to time and accurately delay the expression of lysis proteins by engineered bacteria is of great interest to the synthetic biology field, since it is one of the safety mechanisms that is used for bio-containment [33]. Therefore, we wanted to explore the circuit behavior when producing a lysis protein rather than YFP, so a designed circuit that produces the MS2 lysis protein L was designed (see Fig. 2). Although MS2 bacteriophage is one of the most researched bacteriophages, the mechanisms underlying protein L lysis abilities remain mainly unknown [34]. Similar to the YFP assays setup, HSL was present in the media from T=0 and Ara was added at different growth phases and at different concentrations, and the time for lysis was measured (see Methods) (Fig. 9a and b). When the inducers were added at the EL phase, the time for lysis was 240 min, however, when the inducers were added at later growth phases, there was a gradual significant decrease in the time it took to detect lysis (Fig. 9c). Thus, the induction at later growth phases alters the circuit behavior and leads to a faster lysis. Similarly, when a 10:1 concentration was added, the bacteria were lysed faster than the bacteria cultivated with a 1:1 concentration (Fig. 9d).

**Figure 9:**
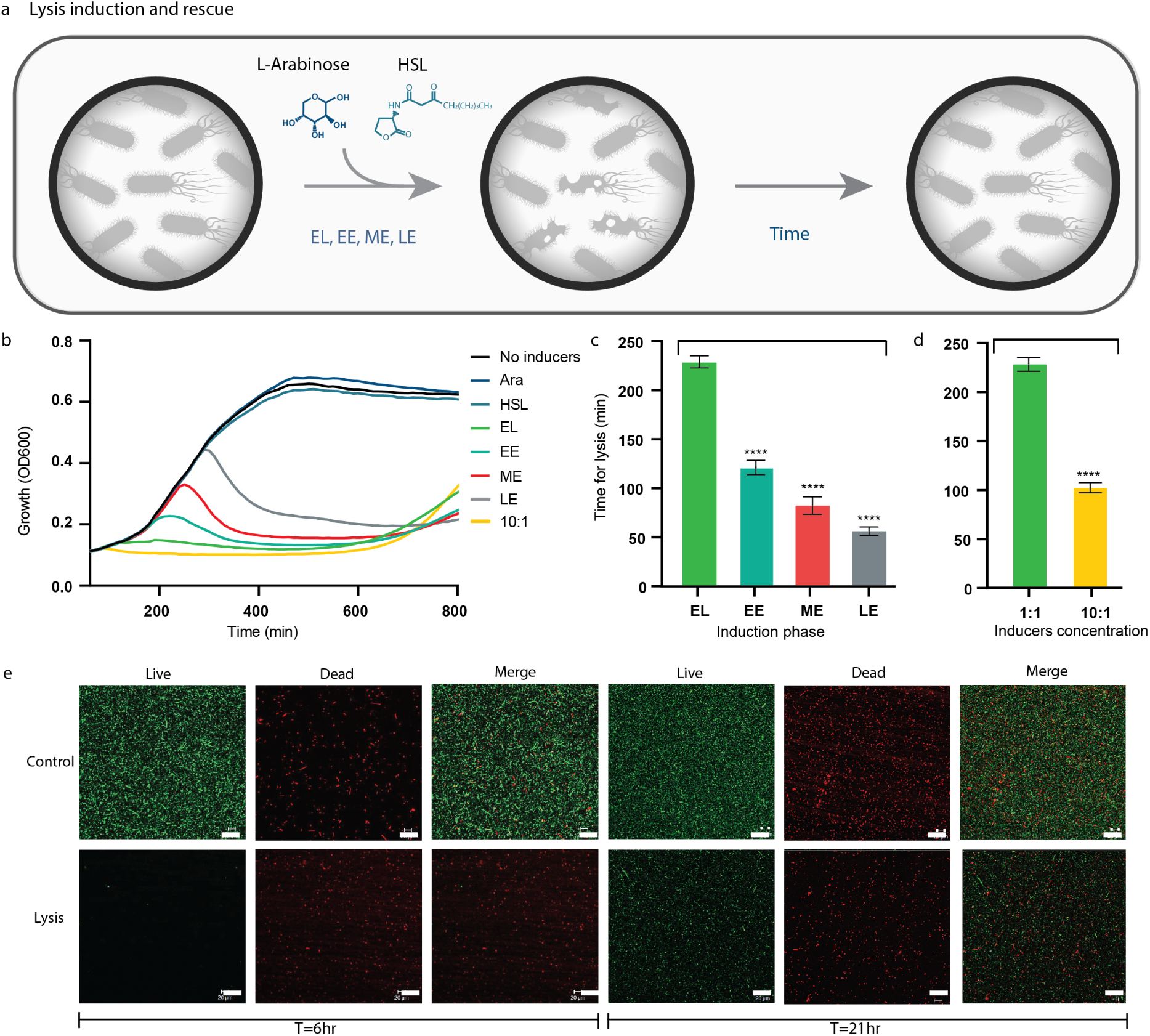
Lysis induction under different conditions. **a** Scheme of the lysis assay. HSL was added at T=0, Ara was added later at different growth phases which induced lysis. With time, some bacteria overcame the lysis process, which is shown with OD. **b** Bacteria growth over time of the different bacteria induced at various growth phases. **c** and **d** Comparison of the time it took to achieve a decrease in the OD of the bacteria at different growth phases (c) and inducers concentrations (d). \*\*\*\**P* < 0.001 (Student’s *t*-test). **e** Confocal images of a live-dead assay of control bacteria (without induction) and lysed bacteria at two different time points. Scale bar is 40 *μ*m.

The OD was measured for 800 min, and around 650 min there was an increase in the OD across all samples which led us to believe that the bacteria escaped the lysis process, probably due to mutation insertions in the circuit plasmids. It should be noted, that the time it took the bacteria to rescue itself from lysis was the highest for EL induction (T=0) and the 10:1 inducers’ concentrations (also at T=0) (Supplementary Fig. 3). This can be explained by the initial small amount of bacteria at T=0. Samples from different time points were taken to a confocal microscopy for a live-dead assay and compared to a non-induced sample (Fig. 9e). The results show that indeed following 6 hours from induction. The vast majority of bacteria were dead, however, following 21 hrs, there was an increase in the amount of live bacteria, further validating the ability of the bacteria to escape lysis (Fig. 9b and e). These results indicate that for an efficient bio-containment control, the L protein of the MS2 bacteriophage should be combined with an additional killing mechanism, regardless of the different induction conditions.

## 3 Discussion

The DBTS cycle is a powerful methodology that is commonly and successfully implemented in the synthetic biology field. Yet, it is not flawless and is constantly needing to be re-examined to reduce the turnaround of synthetic biology applications. As an example, when engineering bacteria to act as a sensor for a specific molecule, the two features of time for fluorescence detection and signal intensity are very important. This work shows that if bacteria was sensing a molecule at the *EE* stage, and was observed prior to its ability to produce a fluorescence signal, then it can lead to a false negative result. In addition, bacteria that sensed the molecule at the *S* phase, and therefore produced a significantly low fluorescence signal, can also lead to a false negative result. Furthermore, our results support the notion that when designing a genetic circuit, the range of inducer concentrations that can lead to a satisfying performance can vary across different cultivation temperatures which is a major factor when transitioning outside the lab. Additionally, in this work we hypothesized the values of 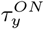 shown in Fig. 8a vary for the different growth phases of the clonal bacteria. After learning from the appropriate test results, we could acquire new models in the *Scale* step that could predict the behavior of these circuits on untested experimental conditions. Thus, by introducing the *Scale* step and broadening the *Test* step to include more environmental factors will subsequently advance the overall learning and will provide deeper understanding of the obstacles that genetic circuit’s performance face when taken outside of the lab.

Expansion of the *Test* step will inevitably promote new prediction models and tools developed in the *Scale* step, able to predict the behavior of genetic circuits under different conditions (even untested ones), and therefore will enable better design choices than those provided by GDA tools. Furthermore, these studies can identify, which experimental conditions have a greater effect on a genetic circuit’s performance. Currently, there is a lack of standardized methodologies and/or software tools to help researchers perform a meaningful *Scale* step and benefit from its results. Even popular GDA tools (like Cello [20]), which provide extensive and automated *Design, Build*, and even *Test* steps, lack of a proper *Scale* step, which will be beneficial to help researchers with better design choices for OTL genetic circuit applications. However, as of now, there is no consensus on what these methodologies should look like, nor tools to help with this process.

## 4 Methods

### 4.1 Circuit Design

A naive implementation of delay in a genetic circuit is shown in Fig. 10a, where successive pairs of NOT gates can be used to add delay to a circuit (from a change of inputs to a change in outputs) without changing the circuits’ function or behavior. However, this circuit can produce unwanted output production (*set-up* glitches) when it is initialized, even in the absence of input molecules, since its components have not been stabilized yet [35]. This means that when the circuit is initialized, since the circuit is not stabilized, some internal gates will start randomly producing output before others. This can cause an erroneous or faulty initial state for the circuit, and therefore an unexpected or unwanted output protein production. So, for example, the gate that produces the output protein of a circuit (blue gate, Fig. 10a) could start producing output before the circuit reaches steady-state without any inducer (input) present, at which point it will be repressed by the green gate.

**Figure 10:**
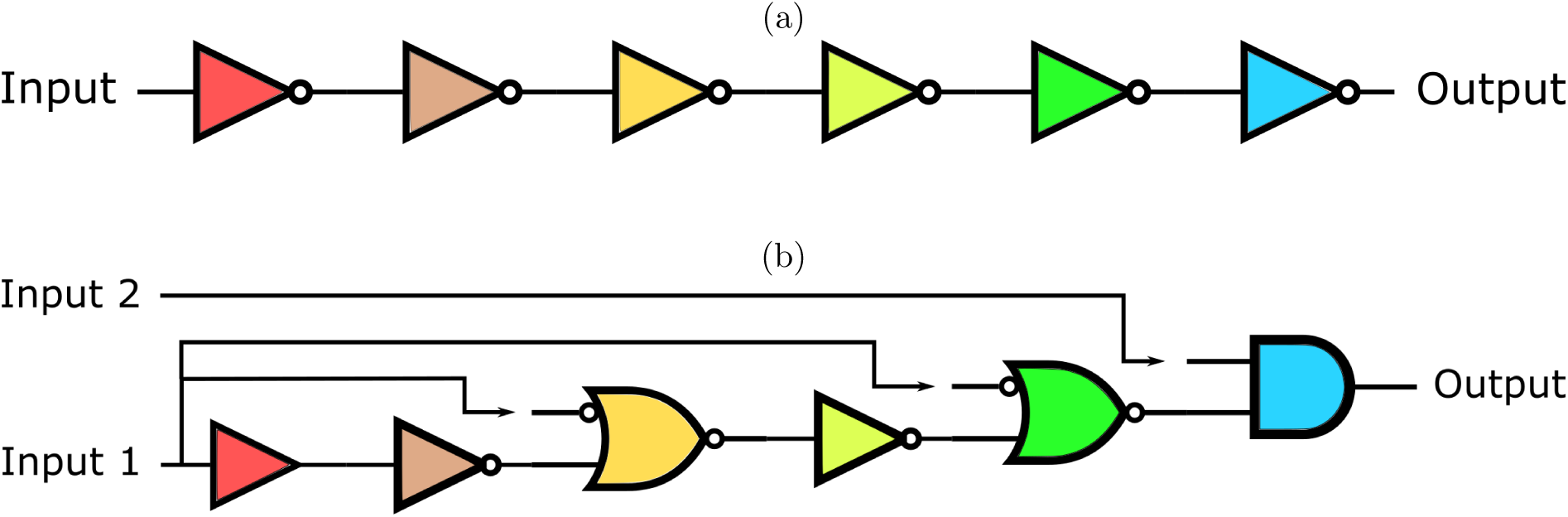
(a) Simple delay circuit. Two successive NOT gates (represented as 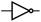) add delay to a circuit without changing the circuit’s behavior. In this image, each logic gate is represented with a different color to represent different gate assignments. (b) Set-up failure-free delay circuit. In this figure 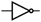 represents a NOT gate, 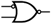 an NIMPLY gate, and 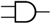 AND gate, and 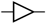 a *buffer* gate. Each logic gate is represented with a different color to represent different gate assignments.

However, it is possible to re-design the circuit in a way to avoid these initialization problems and properly locking the initial-state down, so that there is no unwanted switching behavior, or set-up glitches, when this circuit is initialized. Fig. 10b shows such a design that would avoid set-up failures due to the initialization problem. When such a circuit is transformed into a bacteria, and there is random production from internal gates since the circuit is not in steady-state, there will be no unwanted output production. This is because the output-producing gate is an AND gate, which needs the presence of two signals before it can produce the output signal. The second inducer is necessary so that even if there is some initial leakage production of the green gate, the output is not going to be produced. The results section shows the implementation of Fig. 10b using Cello gates [20].

### 4.2 Mathematical Model

The model used by the automatic model generator of this work is based on a combination of a steady-state model developed in Nielsen et al. [20] and Shin et al. [31], with a dynamic model developed by Moser et al. [36]. The modeling and simulation in this work uses Cello genetic parts and parametrization, but it can work for any genetic circuit as long as the appropriate parameters are available.

The mathematical model used in this work is explained and implemented in Fontanarrosa et al. [35]. However, certain adaptations have been made to this model to be able to account for a “split” sensor gate in the design of the delay circuit. The model developed in Shin et al. [31] represent sensor and internal genetic gates differently: while internal gates are modeled from a Hill-function-like equation, sensor gates are modeled as either being *ON* or *OFF*. Therefore, we adapted the model in order to have a Hill-function-like equation for sensor gates too, in order to model the circuit shown in Fig. 10b. This circuit’s second-to-last gate (green gate), is a sensor gate, which was not initially intended to be an internal gate [20].

The model uses response functions to describe the steady-state RNAP flux output (in RPUs) of a gate over the *output promoter* (promoter that the gate has an effect on), as a function of the RNAP fluxes of the input promoters (for greater detail, see [35]). However, for the model developed in Shin et al. [31], the steady-state calculation of input (sensor) promoter activities of input gates takes the following form:

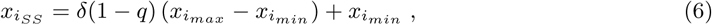

where 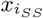 is the steady-state output RNAP flux of sensor gate 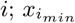 and 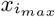 are the minimal and maximal output RNAP fluxes, respectively, for sensor gate *i*; *q* is the presence (*q* = 1) or absence (*q* = 0) of inducer molecules; and *δ*(1 *− q*) is 1 when there are inducer molecules present and 0 when there are not. This formula shows that the response function for a sensor gate is *digital* : it is either *ON* or *OFF*, there is no response curve. This formulation would not work if a sensor gate is used as an *internal* gate as is used in the designed circuit of this work. Therefore, an adaptation of equation 6 to emulate an internal gate response function model was implemented in this work.

The mathematical models for the internal gates are also taken from Shin et al. [31], in which they describe the dynamical behavior of these gates. Equation 7 describes the steady-state output of the internal gate as:

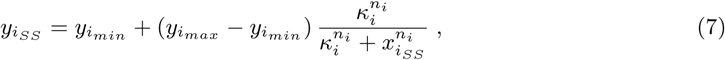

where 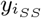 is the steady-state output RNAP flux of gate 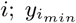 and 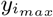 are the minimal and maximal output RNAP fluxes, respectively, for gate *i*; *κ*_*i*_ and *n*_*i*_ are obtained from the affinity and cooperativity of transcription factor binding; and, finally, 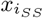 is the input RNAP flux from the input promoter calculated in equation 6. This would give the steady-state output of the gate used to calculate the dynamic behavior of it.

However, to describe the dynamic behavior of the internal gates, a set of ODEs for each genetic gate is needed to describe the timescale by which a gate turns *ON* or *OFF*, using a simplified model that uses only two parameters (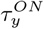 and 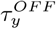) as shown in the following equation:

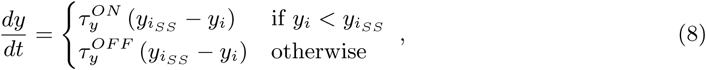

where 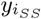 is the RNAP flux of gate *i* at steady state (Equation 7), *y*_*i*_ is the current RNAP flux of gate *i*, and 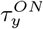 and 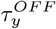 which are the bundled kinetic parameters that capture the response time to go to a steady state that is higher than the current output 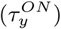 or lower 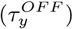 [31, 36], and were set to an average value [31] (see supporting information). Finally, to calculate the RPU output of the promoter controlling YFP expression we use Equation 9:

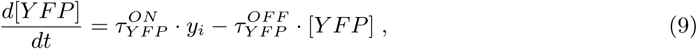

where 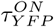 and 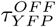 which are bundled kinetic parameters that capture the response time to go to a steady state that is higher than the current output 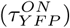 or lower 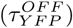 [31, 36], *y*_*i*_ is the non-additive input RNAP flux of the previous internal gate, and [*Y FP*] is the fluorescence output measuring YFP expression.

The values for the parameters of the first non-informed model predictions were taken from literature [31], or used averages for when there were no values for certain gates (i.e. for sensor gates used as internal gates). However, after the re-characterization of the parts, these parameter values where changed to use the parametrization values (see results section). The resulting complete model is then analyzed using the Runge-Kutta-Fehlberg (4,5) method [26] implemented in iBioSim [25], using the Synthetic Biology Open Language [37] to describe the design. The model and parameter values, can be seen in the Supplementary Information.

### 4.3 Parametrization Methods

#### Hill-function characterization algorithm

For the Hill-function parametrization method, a normalized least-squares method using the non-linear *Least-Squares Minimization and Curve-Fitting* (lmfit) Python package [32] was used, and random initial parameter estimations following the GAMES workflow [38].

#### ON/OFF and OFF/ON characterization algorithm

The lmfit Python package, which is based on the Levenberg-Marquardt minimization algorithm, was used to perform the fits and analyze the resulting parameter sets [32]. The fits were performed by minimizing the sum of the square of the relative error between each measured data point and the same point in a corresponding model simulation. As with the Hill-function characterization algorithm a random initial parameter value search was implemented following the GAMES workflow [38], while simultaneously looking for the smallest chi-squared values for each fitting iteration. These scripts are in the supplementary information documentation.

Using the estimated values of 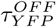, shown in Table 4, and using both Equations 3 and 4, the first fitting iteration was used to obtain 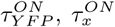, and *x*_*ss*_ parameter estimate values using the *ON-to-OFF* characterization experiment results. Using the parameter estimation method proposed in [38], and the fixed values of 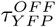 obtained previously, the model was fitted to the experimental results using a minimizing function. The parameter values estimated with this method are shown in Table 4.

Using the *ON-to-OFF* characterization experiments, and assuming that the influence of input sensor promoter flux is zero, then fitting Equation 4 to the gradient of the fluorescence loss over time produces estimates of 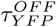 parameter values.

### 4.4 Plasmids preparation

#### Circuit plasmid (SZT61)

pAN3944 containing the AraC gate sequence, pAJM.477 containing the Lux R sequence and pAN4023 containing the Lux R relevant ribozyme (RiboJ), RBS (BBa B0064 rbs) and terminator (L3S2P21 terminator) were a gift from Prof. Christopher Voigt (Addgene plasmids #74702, #108526 and #74701, accordingly). The Lux R sequence from pAJM.477 was amplified using *polymerase chain reaction* (PCR) and cloned into the relevant location at pAN4023 using a standard Gibson assembly reaction [39]. Then, the entire Lux R gate was amplified using PCR and cloned into pAN3944 using standard Gibson assembly reaction. The removal of other gates from pAN3944 was done using one-step PCR.

#### plasmid containing YFP (SZT45)

Using one-step PCR, the promoter of the reporter plasmid pAN4023 was changed to pLux Star.

#### Reporter plasmid containing the MS2 lysis protein (SZT65)

The lysis protein sequence was synthesized (genscript) and cloned into SZT45 using standard Gibson assembly reaction s[39].

#### Lux R gate characterization circuit plasmid (on/off) (SZT69)

AraC gate was removed from SZT61 and the LuxR promoter was changed from the pBAD to a constitutive promoter - J23105, using reverse PCR.

#### AraC gate characterization circuit plasmid (on/off) (SZT70)

LuxR gate was removed from SZT61 using reverse PCR.

#### AraC gate characterization reporter plasmid (on/off) (SZT71)

The pBAD promoter was amplified from pAN3944 and cloned into SZT45 instead of pLuxStar promoter using standard Gibson assembly reaction.

### 4.5 Circuit Induction and Measurements

The genetic circuit plasmid (SZT61) and the relevant reporter plasmid (SZT45 or SZT65) were cotransformed into chemically competent NEB 10-beta (New England Biolabs, MA, C3019) according to the manufacture instructions. Following the transformation, the cells were plated on LB agar plates with 50 *μ*g/mL kanamycin (Gold Biotechnology, MO, K-120-5) and 50 *μ*g/mL spectinomycin (Gold Biotechnology, MO, S-140-5). The plates were grown at 37°C overnight and single colonies were chosen and inoculated into 200 *μ*l of M9 glucose with antibiotics in a deep 96-well plate (MasterBlock, 96 wells, PP, 2ml). M9 glucose media is composed of M9 media salts (6.78 g/L Na2HPO4, 3 g/L KH2PO4, 1 g/L NH4Cl, 0.5 g/L NaCl;), 0.34 g/L thiamine hydrochloride (Sigma-Aldrich, MO, T4625), 0.4% D-glucose (BDH), 0.2% Casamino acids (Bacto), 2 mM MgSO4 (Fisher Chemicals), and 0.1 mM CaCl2. Antibiotic concentrations in M9 glucose media were 50 *μ*g/mL kanamycin and 50 *μ*g/mL spectinomycin. The single colonies were grown at 37°C overnight, 1000 RPM in Multitron Pro 2 shaker incubator. Following the incubation, the overnight cultures were diluted 178-fold by adding 15 *μ*l of the culture into 185 *μ*L of M9 glucose media, and then 15 *μ*L of that dilution into 185 *μ*L of M9 glucose media with 50 *μ*g/mL kanamycin and 50 *μ*g/mL spectinomycin. For the different growth phases, the M9 media contained 2 *μ*M of N-Hexanoyl-L-homoserine lactone (HSL) (Sigma-Aldrich) and the L-arabinose (Ara) (Sigma-Aldrich) was added at the relevant times to a final concentration of 5 mM. For the different concentrations of inducers assays, the M9 media contained the following inducer’s concentrations shown in Table 1:

**Table 1:** The different inducer concentrations that were tested.

| Condition | HSL | Ara |
| --- | --- | --- |
| 1:100 | 0.02μM | 0.05mM |
| 1:10 | 0.2μM | 0.5mM |
| 1:1 | 2μM | 5mM |
| 10:1 | 20μM | 5mM |

For the soil assays, the soil sample was split to two controls, sterile and non-sterile. The sterile soil was autoclaved and the non-sterile soil was not. Then, the two controls were mixed with the M9 media to a final concentration of 2% (W/V). The bacteria were grown according to the description above at M9 media and were diluted 178-fold into the soil media.

#### Fluorescence Measurements

The diluted culture was plated in a black 96-wells plate with a clear bottom (655090,F-bottom, *μ*clear, black, Greiner) and placed in a plate reader (Tekan SPARK plate reader) at 37°C and 270RPM. OD600 and fluorescence (excitation wavelength 485nm and emission wavelength 535nm) were measured every 10 minutes for at least 800 minutes. For the lysis protein assay, only OD600 was measured. For the soil assay, only fluorescence was measured. For the different temperature assays the plate reader temperature was set to 30°C and 42°C

#### ON/OFF Characterization Assays

For the AraC gate, AraC plasmid (SZT70) and the reporter plasmid (SZT71) were co-transformed into chemically competent NEB 10-beta (New England Biolabs, MA, C3019) according to the manufacture instructions. For the LuxR gate, LuxR plasmid (SZT69) and the reporter plasmid (SZT45) were co-transformed into chemically competent NEB 10-beta (New England Biolabs, MA, C3019) according to the manufacture instructions. Following the transformation the cells were plated on LB agar plates with 50 *μ*g/mL kanamycin (Gold Biotechnology, MO, K-120-5) and 50 *μ*g/mL spectinomycin (Gold Biotechnology, MO, S-140-5). The plates were grown at 37°C overnight and single colonies were chosen and inoculated into 200 *μ*l of M9 glucose with antibiotics in a deep 96-well plate (MasterBlock, 96 wells, PP, 2ml). M9 glucose media is composed of M9 media salts (6.78 g/L Na2HPO4, 3 g/L KH2PO4, 1 g/L NH4Cl, 0.5 g/L NaCl;), 0.34 g/L thiamine hydrochloride (Sigma-Aldrich, MO, T4625), 0.4% D-glucose (BDH), 0.2% Casamino acids (Bacto), 2 mM MgSO4 (Fisher Chemicals), and 0.1 mM CaCl2. Antibiotic concentrations in M9 glucose media were 50 *μ*g/mL kanamycin and 50 *μ*g/mL spectinomycin. The single colonies were grown at 37°C overnight, 1000 RPM in Multitron Pro 2 shaker incuabtor. Following the incubation, the overnight cultures were diluted 178-fold by adding 15 *μ*l of the culture into 185ul of M9 glucose media, and then 15 *μ*L of that dilution into 185 *μ*L of M9 glucose media with 50 *μ*g/mL kanamycin and 50 *μ*g/mL spectinomycin. For the AraC gate, the L-arabinose (Ara) (Sigma-Aldrich) was added at the relevant times to a final concentration of 5mM. For the LuxR gate, HSL was added at the relevant times to a final concentration of 2 *μ*M. The diluted culture was plated in a black 96-wells plate with a clear bottom (655090,F-bottom, *μ*clear, black, Greiner) and placed in a plate reader (Tekan SPARK plate reader) at 37°C and 270RPM. OD600 and fluorescence (excitation wavelength 485 nm and emission wavelength 535 nm) were measured every 10 minutes for 650 minutes. Following 650 minutes, the plate were centrifuged for 2 minutes at 4000RPM and the media was removed from each well. Then, fresh M9 media without any inducers was added to all the wells. The plate were then placed in the plate reader at 37°C and 270RPM. OD600 and fluorescence were measured every 10 minutes for an additional 650 minutes.

### 4.6 Circuit Assay’s Analysis

For each sample, there were at least five biological repeats.

#### Fluorescence graphs

The graphs represent the average values of these repeats. The fluorescence was normalized by subtracting the averaged blank value from the averaged fluorescence value and dividing the resulted fluorescence value in the averaged OD600 value for each time point.

#### Time for fluorescence detection graphs

The normalized fluorescence values of the samples from T=0 onward were compared to the normalized fluorescence values of the negative control (without induction). The time of fluorescence detection was determined as the time when the fluorescence values of the samples exceeded those of the negative control.

#### Fluorescence fold change graphs

The maximum normalized fluorescence values of each sample were chosen. For the induction time variations assay, the T=0 average was set as one and all the other samples were compared to it. For the inducer concentration variations assay, the 1:1 concentration average was set as one and all the other samples were compared to it. For the inducer concentration variations at low and high temperature, the 1:1 concentration average was set as one and all the other samples were compared to it.

#### Time for lysis graphs

The normalized OD600 values of the samples from T=0 onward were compared to the normalized OD600 values of the negative control (without induction). The time of lysis was determined as the time when the OD600 values of the samples decreased in comparison to the negative control.

#### Time for rescue graphs

The normalized OD600 values of the samples from the time of lysis were examined. The time of rescue was determined as the time when the OD600 values of the samples started to increase rather than decrease.

#### YFP a.u. production rate graphs

The sum of the normalized fluorescence values of each hour was calculated from the time of the fluorescence signal detection (see time for fluorescence detection graphs explanation).

#### Doubling time graphs

The doubling time was determined as the time (min) it took for normalized OD600 values to double (from 0.2 to 0.4).

#### Confocal images

The bacteria were cultivated and induced as described above for the lysis assay. At T=6hr and T=21hr, 200 *μ*L from each sample was taken and centrifuged at 4000 RPM for 1 minute. The cells were washed using 1 mL PBS and centrifuged. The pellet was resuspended with 50 *μ*L of PBS. Then, a Live/Dead staining was done according to the manufacturer’s instructions, (L13152 LIVE/DEAD® BacLight™ Bacterial Viability Kit, Molecular Probes, OR, USA), and the samples were subjected to confocal microscopy utilizing a LSM510 confocal microscope (Zeiss). The presented results are representative of three independent experiments.

## 6 Author contributions statement

S.Z.T. designed and performed the experiments; P.F. modeled and simulated the different designs, using default and re-characterized parameter values; D.B helped with the live-dead confocal experiments. All authors contributed to conceiving the project, analyzing data and writing the manuscript.

## 7 Acknowledgments

S.Z.T was supported by the Marian Gertner Institute for Medical Nanosystems. The authors thank to Dr. Alex Barbul for help with the confocal images and members of the Gazit laboratory and the Myers laboratory for helpful discussions.The authors also thank the developers of lmfit [32] for making their code available on a free and open-source basis. The authors P.F. and C.J.M. of this work were supported by DARPA FA8750-17-C-0229. Any opinions, findings, and conclusions or recommendations expressed in this material are those of the author(s) and do not necessarily reflect the views of the funding agencies.

## Footnotes

1 https://synbiohub.programmingbiology.org/public/Eco1C1G1T1/Eco1C1G1T1_collection/1

