## Supplementary for "Investigating and Modeling the Factors that Affect Genetic Circuit Performance"

### Supplementary Information: Investigating And Modeling the Factors that Effects the Performance of Genetic Circuits

May 16, 2022

### Parameter Values Estimations

The parameters values used in this work's model predictions are described in this section. The values for the parameters of the first non-informed model predictions were taken from literature [1], stored in SynBioHub. or used averages for when there were no values for certain gates (i.e. for sensor gates used as internal gates). The parameter values used for the rest of the model predictions were obtained from a fitting algorithm using the lmfit Python package [2]. The data sources and scripts are available at a Github repository.

#### Non-informed default parameter values

##### Response function parameters

Table 1: Input Sensor Gate Parameters. This table summarizes the parameters used to calculate input promoter activities in the model.

| Gate | $x_{min}$ [1] | $x_{max}$ [1] |
|---|---|---|
| Tac | 0.0034 | 2.8 |
| Tet | 0.0013 | 4.4 |
| BAD | 0.0082 | 2.5 |

Table 2: Gate Parameters. This table summarizes the parameters used to determine promoter activities of the circuit.

| Gate | $y_{min}$ [1] | $y_{max}$ [1] | $\kappa$ [1] | $n$ [2] |
|---|---|---|---|---|
| AmtR | 0.06 | 3.8 | 0.07 | 1.6 |
| BetI | 0.07 | 3.8 | 0.41 | 2.4 |
| BM3R1 | 0.01 | 0.8 | 0.26 | 3.4 |
| HlyIIR | 0.07 | 2.5 | 0.19 | 2.6 |
| PhlF | 0.02 | 4.1 | 0.13 | 3.9 |
| SrpR | 0.007 | 2.1 | 0.1 | 2.8 |

##### Gate dynamics

We used generic values for $\tau_{ON}$ and $\tau_{OFF}$ parameters obtained from [1]. The values used in this work are $\tau_{ON} = 1.7\ auRPU^{-1}hr^{-1}$ and $\tau_{OFF} = 4.044444444\ auRPU^{-1}hr^{-1}$.

#### Parameter Value Estimations

##### Hill Function Parameter Value Estimations

##### Dynamic Parameters ($\tau_{ON}$ and $\tau_{OFF}$) Value Estimations

### Doubling time

Doubling time across different assays.

### Lysis Protein Circuit Design

### Lysis rescue

Time for rescue of bacteria induced at different growth phases and with different inducers' concentrations.

[1] In units of RPU [3].
[2] Dimensionless.

Draft

Table 3: Hill-function parameter value estimations for different gates, obtained by fitting Equations 1 and 2 to part-characterization experiments at different growth-phases (EL: early-lag, LE: late-exponential). Values rounded to three significant digits.

| | AraC | | | |
|---|---|---|---|---|
| Growth-phase | $y_{max}$ | $y_{min}$ | $\kappa$ | $n$ |
| EL | 1730 | 342 | $4.16 \times 10^{-2}$ | 2.60 |
| LE | 1150 | 366 | $6.68 \times 10^{-2}$ | 2.89 |
| | LuxR | | | |
| EL | 4310 | 223 | 2.33 | $8.40 \times 10^{-1}$ |
| LE | 3230 | 308 | $5.52 \times 10^{-1}$ | $8.08 \times 10^{-1}$ |

Table 4: Dynamic parameter value estimations for different gates, obtained by fitting Equations 3 and 4 to *ON-to-OFF* and *OFF-to-ON* part-characterization experiments for different growth-phases (EL: early-lag, LL: late-lag, EE: early-exponential, ME: middle-exponential, LE: late-exponential, S: stationary). Values rounded to three significant digits.

| | AraC | | | |
|---|---|---|---|---|
| Growth-phase | $\tau_x^{ON}$ | $\tau_{YFP}^{ON}$ | $\tau_{YFP}^{OFF}$ | $x_{ss}$ |
| EL | 0.0822 | 0.126 | 0.0893 | 976 |
| LL | 0.169 | 0.109 | 0.0933 | 1000 |
| EE | 0.117 | 0.13 | 0.114 | 1000 |
| ME | 0.111 | 0.112 | 0.111 | 999 |
| LE | 0.275 | 0.0749 | 0.096 | 998 |
| S | 0.208 | 0.0538 | 0.105 | 1000 |
| | LuxR | | | |
| EL | 0.143 | 0.335 | 0.166 | 995 |
| LL | 0.195 | 1.3 | 0.221 | 211 |
| EE | 0.11 | 0.274 | 0.11 | 938 |
| ME | 0.173 | 0.234 | 0.173 | 977 |
| LE | 0.0729 | 0.173 | 0.0726 | 1000 |

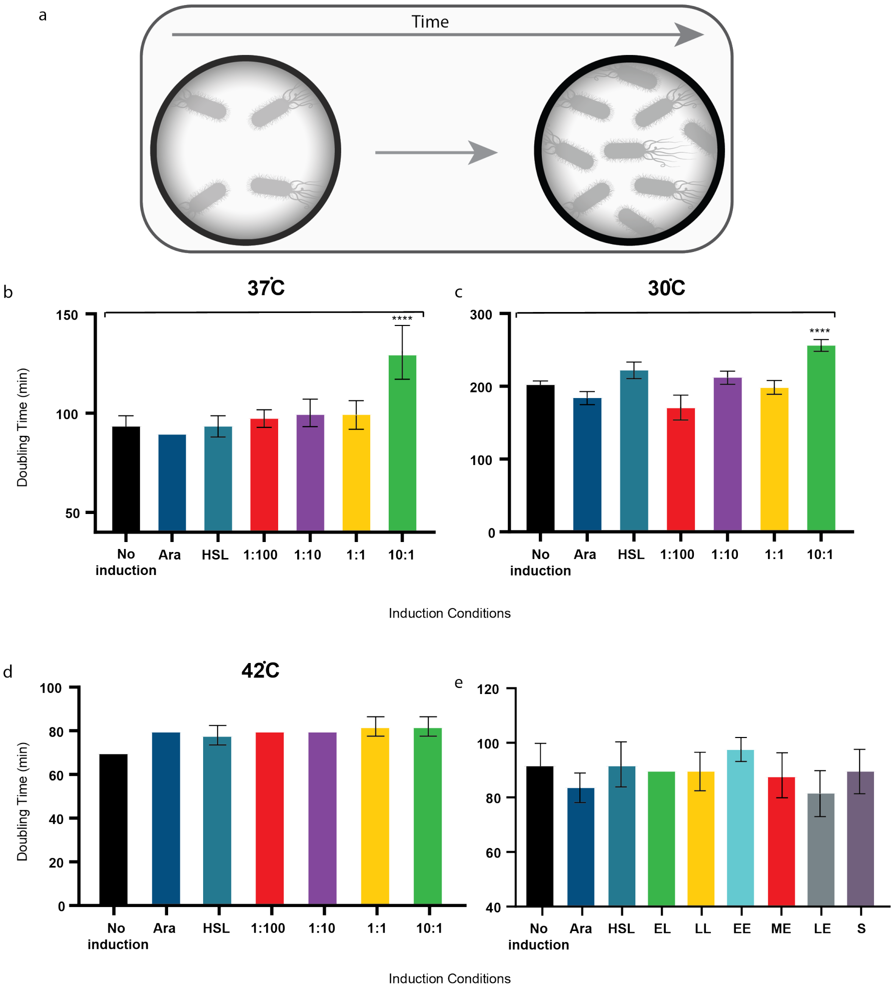

Figure 1: a. A scheme of the doubling time assay, see methods. b-d. Comparison between the averaged doubling times of the bacteria induced with a range of inducers' concentrations and at different temperatures: b. 37°C, c. 30°C and d. 42°C. $****P < 0.001$, student $t$-test. e. Comparison between the averaged doubling times of the bacteria induced at different growth phases.

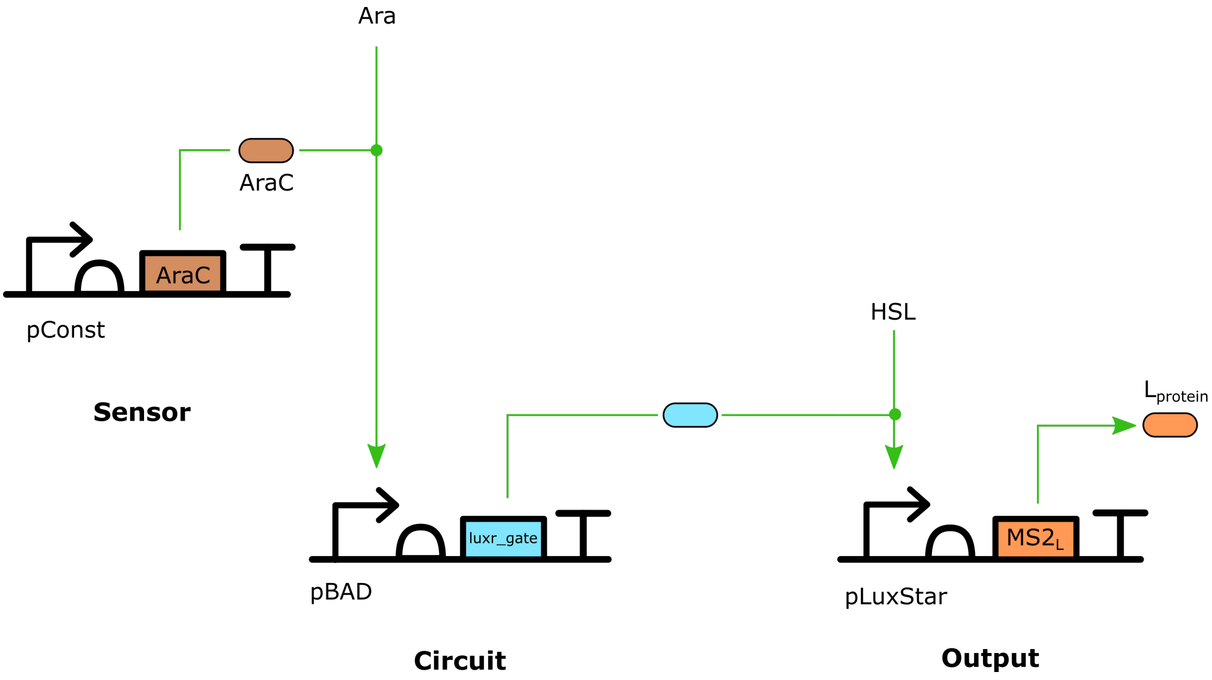

Figure 2: Lysis protein circuit. This circuit is designed to work the same as the delay circuit, but having a lysis protein as an output. This was used to study how changing the output of a circuit might affect the delay of the circuit.

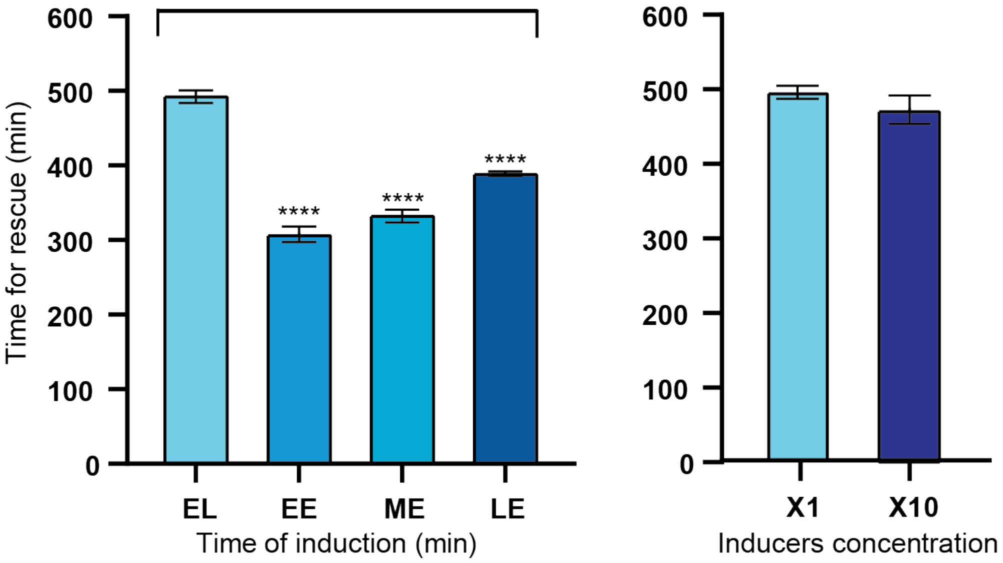

Figure 3: Time for rescue of bacteria induced at different growth phases and with different inducers' concentrations. $^{****}p < 0.0001$ student $t$-test.

#### Vectors map

The different vectors that were used.

##### Delay circuit vectors

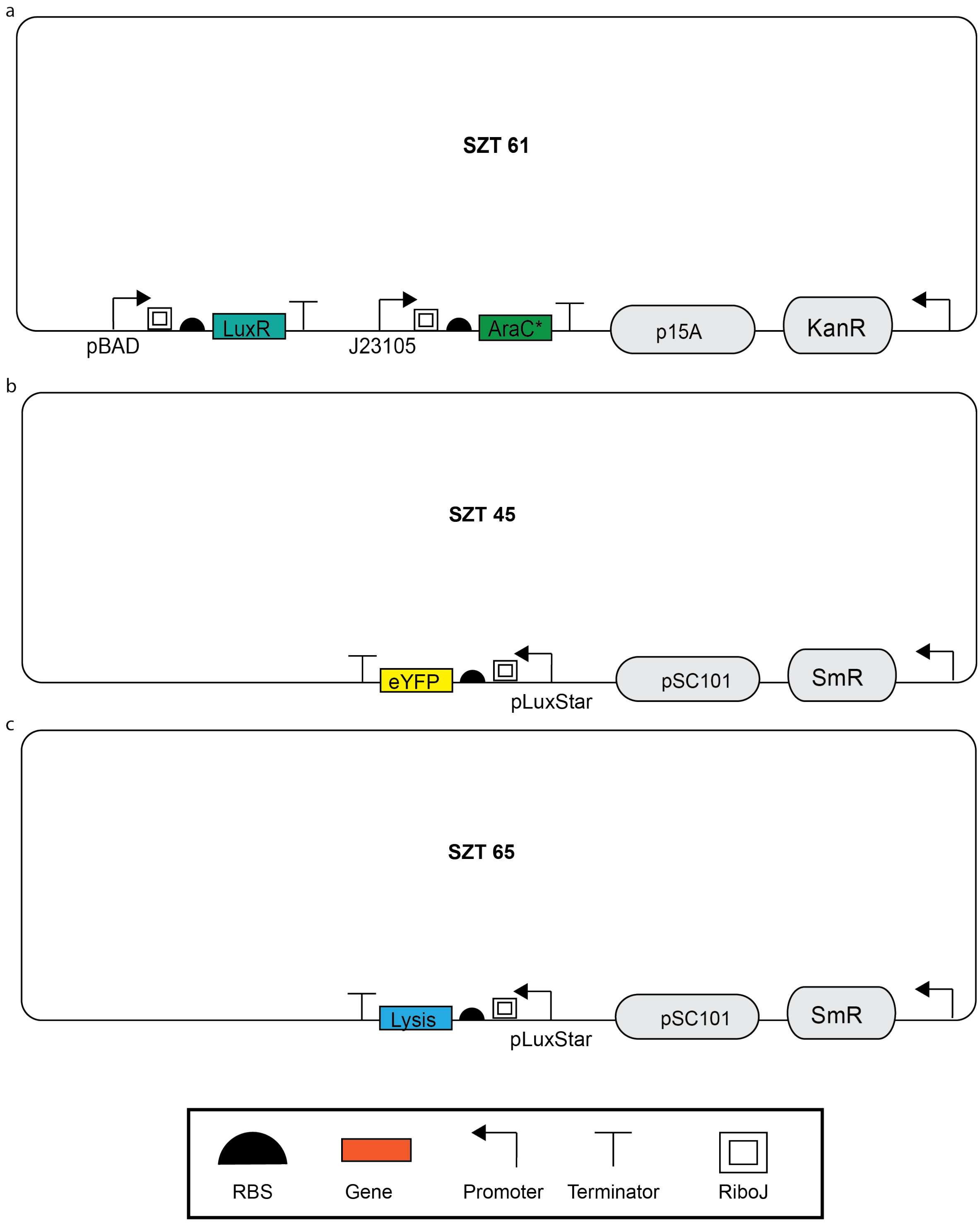

Figure 4: Vector maps of a. delay circuit, b. reporter plasmid encoding YFP, c. reporter plasmid encoding lysis protein.

#### Gates characterization vectors

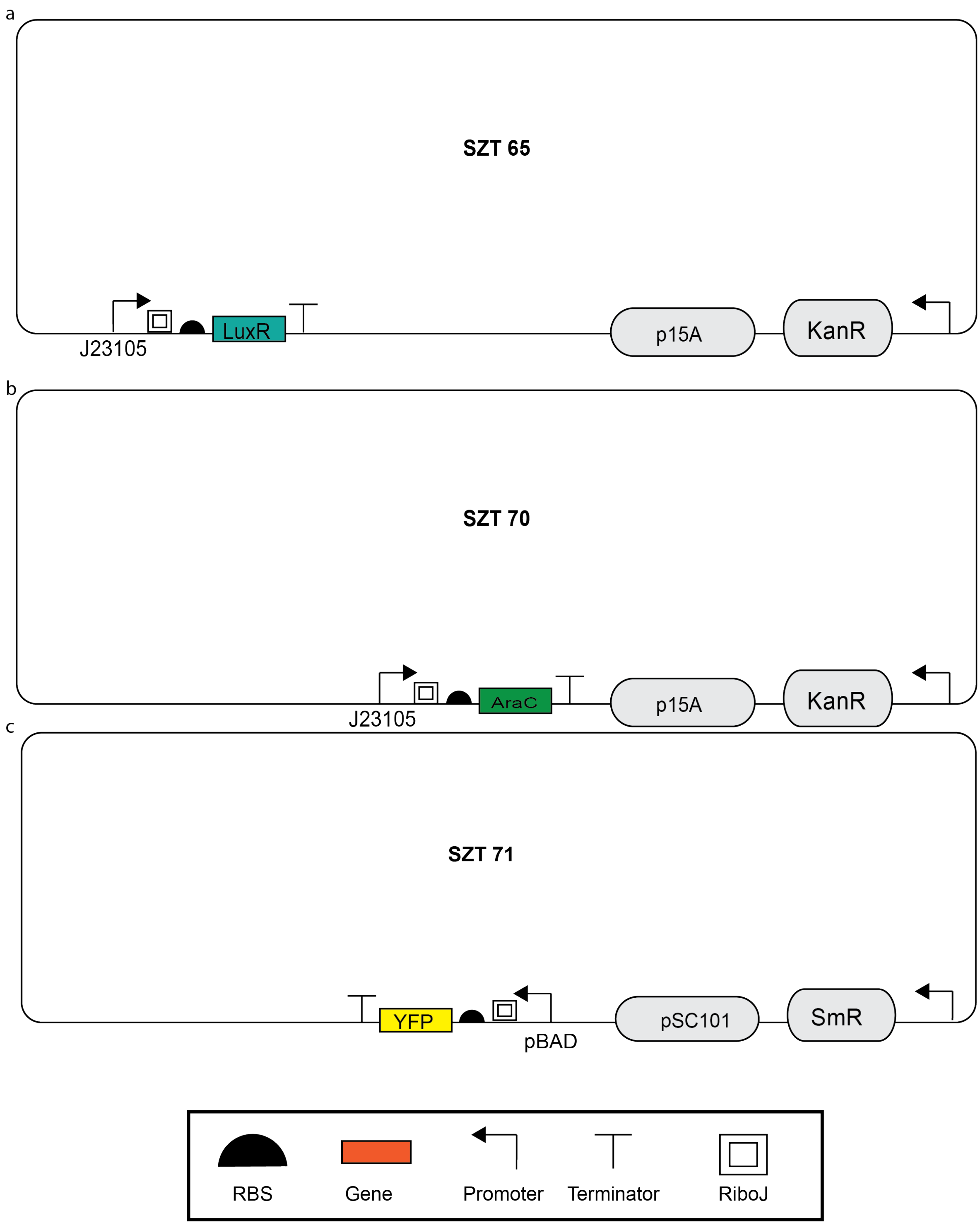

Figure 5: Vector maps of a. LuxR gate characterization, b. AraC gate characterization, c. reporter plasmid for AraC gate characterization.

[2] Matt Newville, Renee Otten, Andrew Nelson, Antonino Ingargiola, Till Stensitzki, Dan Allan, Austin Fox, Faustin Carter, Michał, Ray Osborn, Dima Pustakhod, lneuhaus, Sebastian Weigand, Glenn, Christoph Deil, Mark, Allan L. R. Hansen, Gustavo Pasquevich, Leon Foks, Nicholas Zobrist, Oliver Frost, Alexandre Beelen, Stuermer, azelcer, Andrew Hannum, Anthony Polloreno, Jens Hedegaard Nielsen, Shane Caldwell, Anthony Almarza, and Arun Persaud. Lmfit/lmfit-py: 1.0.3. Zenodo, October 2021.

[3] Jason R Kelly, Adam J Rubin, Joseph H Davis, Caroline M Ajo-Franklin, John Cumbers, Michael J Czar, Kim de Mora, Aaron L Glieberman, Dileep D Monie, and Drew Endy. Measuring the activity of biobrick promoters using an in vivo reference standard. *Journal of biological engineering*, 3(1):1–13, 2009.
